# Cell-type specific repertoire of responses to natural scenes in primate retinal ganglion cells

**DOI:** 10.1101/2025.04.07.647118

**Authors:** Alexandra Kling, Nora Brackbill, Colleen Rhoades, Alex Gogliettino, Alexander Sher, Alan Litke, E.J. Chichilnisky

## Abstract

At least 20 distinct retinal ganglion cells (RGC) types have been identified morphologically in the primate retina, but our understanding of the distinctive visual messages they send to various targets in the brain remains limited. Here, we use large-scale multi-electrode array recordings to examine how multiple functionally-distinct RGC types in the macaque retina respond to flashed natural images. Responses to white noise visual stimulation were used to functionally identify 936 RGCs of 12 types in three recordings. Each cell type was confirmed by the mosaic organization of receptive fields, and 7 cell types were cross-identified between recordings. The average kinetics of light response in each RGC type as well as the repertoire of distinct firing patterns that each type produces were examined across thousands of natural images. The kinetics of the average response across images were highly stereotyped among cells of each cell type and distinct for cells of different types. Moreover, the full repertoires of firing patterns produced by different cell types, assessed by their latency and duration, were generally quite distinct with only a few exceptions. Together these data provide an overview of the range of responses to natural images transmitted from the primate retina to the brain.

## Introduction

At least 20 types of retinal ganglion cells (RGCs) convey visual information to diverse brain regions, including the lateral geniculate nucleus, superior colliculus, and pretectum, where it is further processed to support perception and visually guided behavior ^1,2^. Much progress has been made in characterizing the visual signaling properties of the numerically dominant primate RGCs (ON and OFF parasol and midget cells) using artificial stimuli, such as white noise, contrast steps, and frequency chirps ^3–6^. More recently, the responses of these major cell types to naturalistic images, which contain richer spatial and temporal structure that engages the retina in complex ways, have also been examined ^5–8^, revealing substantial deviations from traditional models of RGC response. However, few studies have examined responses to naturalistic stimuli in the ∼16 or so lower-density RGC types ^9,10^, which constitute about a third of the fibers in the optic nerve and have different retinal connectivity and patterns of projection in the brain than the four numerically dominant types. Thus, our understanding of visual signaling in natural conditions by the diverse visual pathways emanating from the primate retina remains limited.

As a first step, in this study we characterize the average response and response repertoire of diverse primate RGC types to a large set of naturalistic stimuli. By combining large-scale multi-electrode array recordings with quantitative analysis of response dynamics, we show that the temporal neural code is highly varied across RGC types: some pairs of cell types exhibit totally non-overlapping response repertoires, and only a few exhibit substantial overlap. We further show that the diverse naturalistic signaling patterns can be used to distinguish the many RGC types, including cell types not as easily distinguished using traditional stimuli.

## Results

### Kinetics of mean response to natural images vary substantially across cells

The visually evoked spiking activity of hundreds of peripheral primate RGCs was recorded *ex vivo* on a custom 512-electrode recording system ^11,12^. To characterize light response properties and classify cells, a white noise stimulus (flickering checkerboard) was used (see ^3,11,13–15^). Cell types were distinguished based on the spike-triggered average stimulus time courses and the interspike interval distribution observed during white noise stimulation. The accuracy of cell type identification was confirmed by the mosaic organization of the receptive fields of cells of each type ^14^ (Fig. 1).

**Figure 1.**
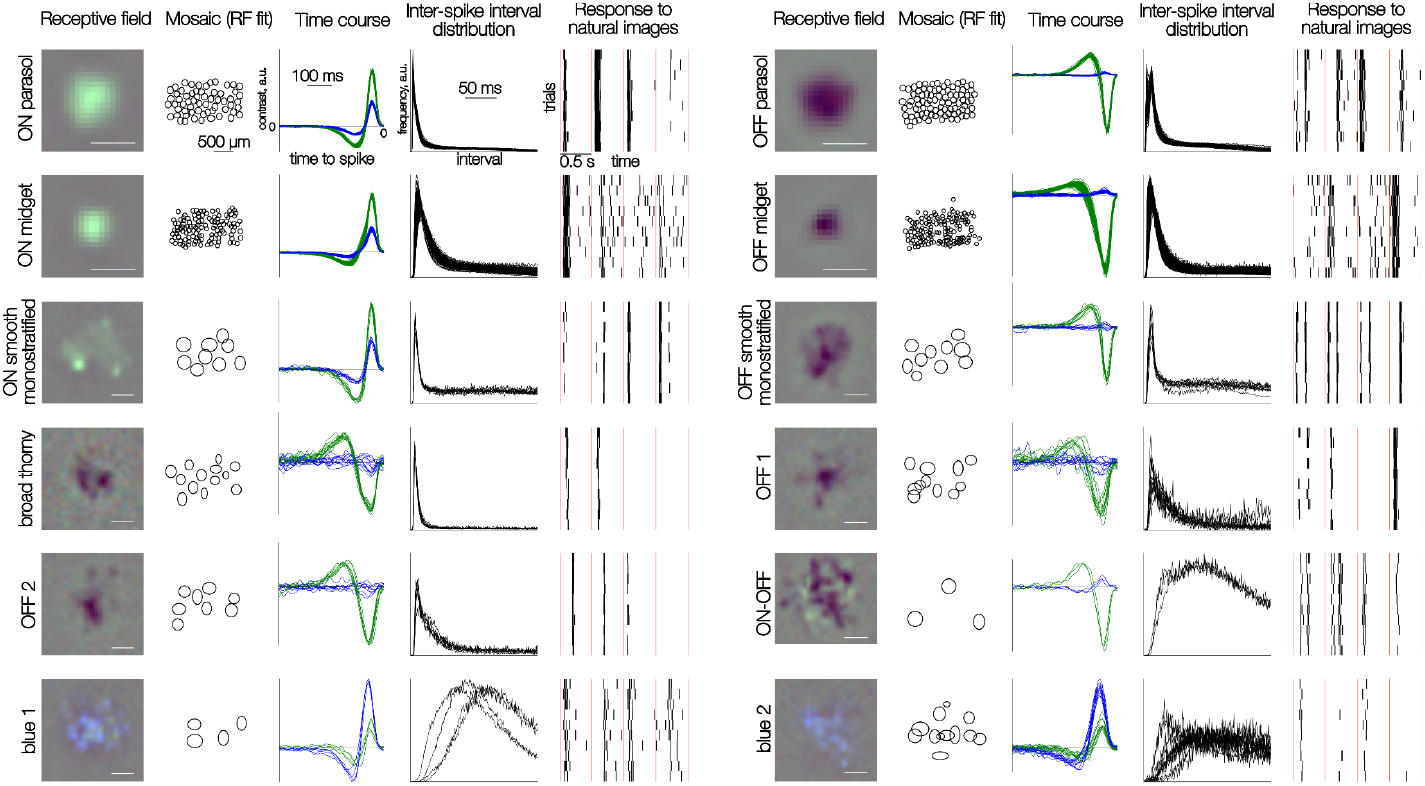
Distinct responses to white noise and natural images in 12 RGC types. Each row of five panels shows results for multiple cells of one type. Left to right: the spatial RF of one cell obtained from the spike-triggered average (see Methods); the mosaic of elliptical fits to the RF profile; time courses for green and blue display primaries obtained from the spike-triggered average; inter-spike interval distributions in the presence of the white noise stimulus; rasters of spike responses to 10 repeated trials of 4 consecutive natural images. Each image was presented for 100 ms; red vertical lines separate distinct image trials (0.5 s trial duration). Scale bar for RFs: 200 µm, for mosaics: 500 µm.

To explore the behavior of the diverse RGC types in naturalistic conditions, the responses to thousands of flashed grayscale naturalistic images were recorded. Each image was presented for 100 ms, followed by 400 ms of a gray background at mean light intensity, a protocol designed to isolate responses to distinct stimuli and to minimize adaptation. A total of 10,000 unique images was shown. A sequence of 150 images was repeated after each block of 1,000 unique images to monitor recording stability. All cells selected for subsequent analysis exhibited reproducible responses to repeated images, with consistent spike patterns in both timing and number (Fig. 1). As expected, the responses of a given cell to distinct images were, in general, markedly different (Fig. 1). Also, as expected, a given image elicited clearly distinct response kinetics in different cells.

To capture the typical kinetics of responses to natural images produced by each cell, its measured responses were aligned to the stimulus onset (Fig. 2a), temporally smoothed, and then averaged across stimuli (Fig. 2b). Despite the variability of responses across images (Fig. 1), the measured firing rate exhibited distinctive kinetics in each cell type (Fig. 2b). Furthermore, the mean response profiles of multiple cells of the same type (identified using white noise stimulus) were highly consistent (Fig. 2b, red lines). This consistency indicates that averaging across a large natural stimulus set extracts a response signature representative of each cell type, not unlike the average signature obtained with white noise stimuli ^14^.

**Figure 2.**
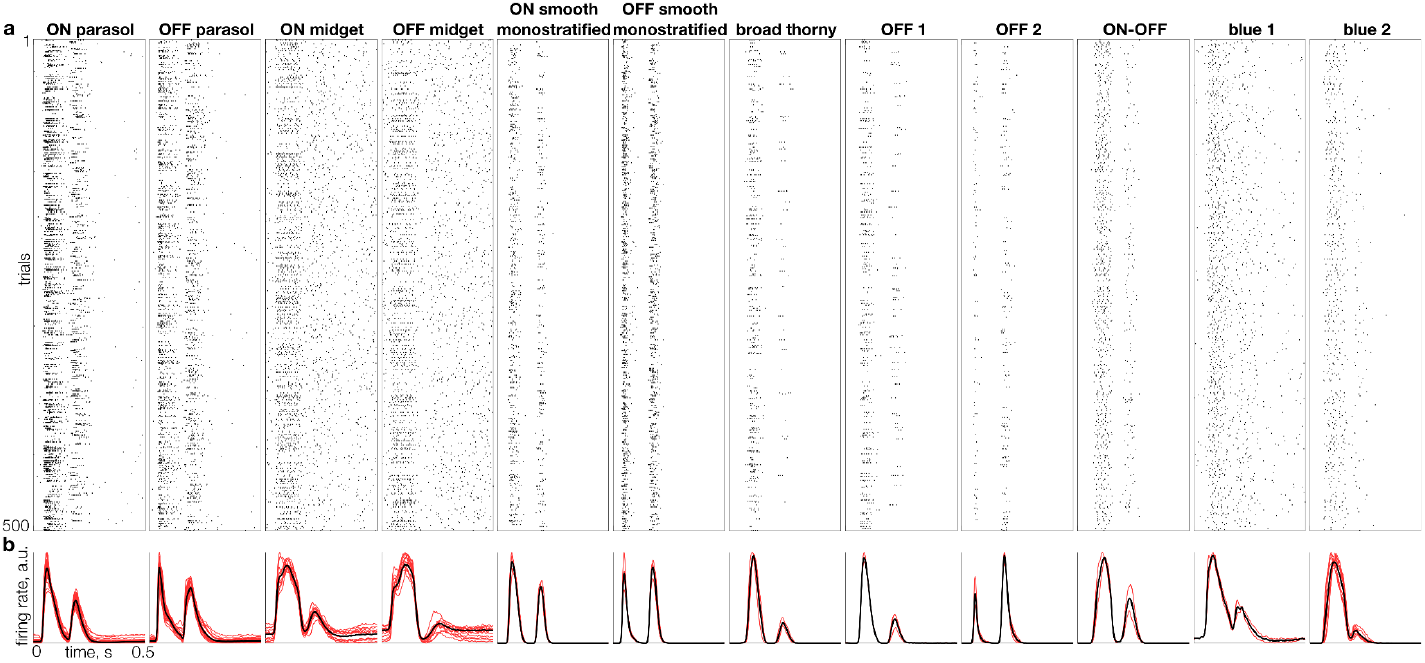
Individual and average responses of 12 RGC types to natural images. **a**. Raster of responses to 500 randomly selected flashed images (rows), for a representative cell of each distinct type (columns). **b**. Mean firing rate across the full set of 10,000 images (L2-normalized) for multiple cells of each type (red traces) and average for all cells of each type (black trace).

Analysis of the kinetics of mean responses to natural images of several cell types revealed several trends. First, the duration of the mean response exhibited a well-known distinction: parasol cell responses are more transient than those of midget cells. Second, for morphologically paired cell types (e.g. ON vs OFF parasol RGCs), OFF cells tended to have a shorter time to peak on average than ON cells (see Fig. 4a), in agreement with previous studies ^16^ (but see ^4^). Third, among low-density cell types, responses were predominantly transient. Interestingly, most of the low-density cell types - ON and OFF smooth monostratified, broad thorny, OFF 1 and OFF 2 types - exhibited more transient responses than parasol cells, while no cell types with more sustained responses than midget cells were identified. Fourth, some cell types - such as parasol and smooth monostratified cells - exhibited strong responses to both image onset and offset (corresponding to the two peaks in Fig. 2b), while other cell types including OFF midget and blue 2 cells responded predominantly to image onset (or more rarely, offset). Together, these observations highlight the diversity of the average retinal code in different cell types.

### C*ell types can be identified by the average kinetics of responses to natural images*

To test if the average response signature could be used to reliably classify RGCs into types, t-distributed Stochastic Neighbor Embedding (t-SNE, ^17^) was applied to the first 10 principal components of the average response profiles of all recorded cells. The results (Fig. 3a) revealed that cells largely cluster into well-defined groups. These groups correspond to the types determined using their responses to white noise stimulation (Fig. 1, 3b). Interestingly, however, the two distinct types of stimuli more clearly and reliably distinguished different sets of RGC types. For example, white noise responses did not reliably discriminate between cells of two OFF-dominated types that had similar inter-spike interval distributions and spike-triggered average time courses (Fig. 3b, black arrows; see Fig. 1, OFF 1 and OFF 2 types) – these cells were separated into two types in the preceding analysis (Fig. 1) largely because they formed two overlapping mosaics ^14^. However, with naturalistic stimuli, these two cell types were readily distinguished (Fig. 3a, black arrows). Conversely, two cell types that produced somewhat similar mean responses to natural images (Fig. 3a, red arrows) were readily distinguished by white noise stimulation (Fig. 3b, red arrows). In fact, a combined t-SNE analysis, incorporating features from both natural image and white noise responses, yielded clearer separation across the complete collection of cell types analyzed (Fig. 3c), indicating that natural image responses provide discriminative power that augments the discriminative power provided by white noise stimuli.

**Figure 3.**
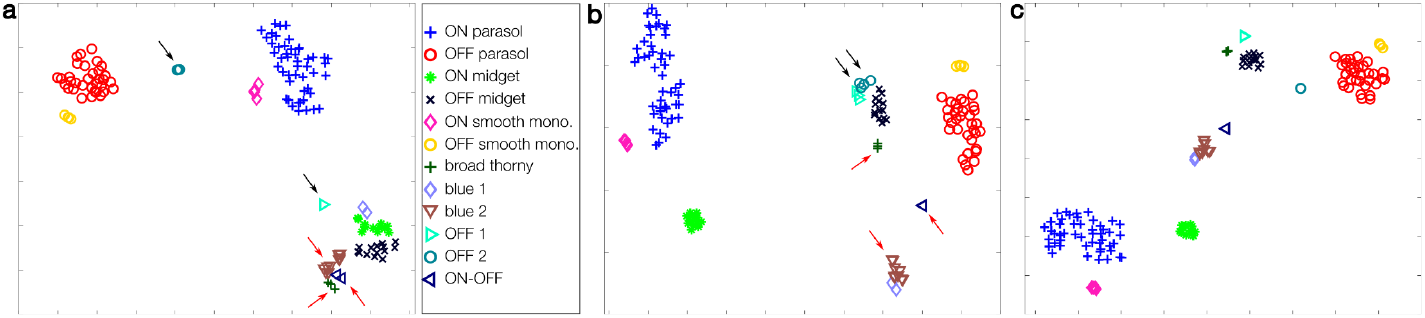
Two-dimensional summary representation of light response properties of RGCs in one recording. **a**. t-SNE representation of recorded firing rates (see Fig. 2) of 141 RGCs in response to flashed natural images. Cell types were identified using responses to white noise stimuli (Fig. 1, see Methods). Black arrows point to well-separated clusters, red arrows point to clusters that are more difficult to distinguish. **b**. t-SNE representation of spike-triggered average (see Methods) responses of the same cells to white noise stimuli. Black and red arrows point to the same groups of cells as in **a. c**. t-SNE for the same cells based on responses to both natural images and white noise.

### Repertoire of responses to natural images is cell-type specific

While the average response kinetics provide a useful summary, they do not reveal the full repertoire of responses produced by each cell to many images. For example, a similar average response in two cells or cell types could be achieved with very different individual responses. Therefore, to understand the similarity of natural image signaling between cells and cell types, a method was developed to compare the repertoire of responses produced by pairs of cells. This comparison was designed to be distinct from comparing the full probability distribution of responses, because in general each cell had a unique receptive field location, and was thus primarily stimulated by a unique set of natural image patches during the experiment.

Specifically, to compare the response repertoires of two cells, the temporal structure of individual responses - latency and duration (see Methods) - was analyzed. While these measures do not fully capture response structure, distinct relationships between cell types were revealed: highly overlapping response repertoires (Fig. 4a), partially overlapping repertoires (Fig. 4b), subset repertoires (Fig. 4c), and fully distinct repertoires (Fig. 4d). In particular, repertoires of cells of the same type typically fully overlapped with each other. Cells of paired ON/OFF types (such as ON and OFF parasol) tended to overlap slightly less, due to slightly shorter latencies in OFF types (Fig. 4b). Interestingly, some cell type pairs revealed complex relationships between their repertoires. For instance, the onset responses of an OFF-dominated cell (OFF 2) formed a subset of the ON parasol cell onset responses, especially in terms of duration, while the collections of offset responses were much more similar (Fig. 4c). Finally, some cell types, such as ON smooth monostratified and broad thorny cells, appeared to have a fully distinct set of latencies - even the slowest responses of a ON smooth monostratified reach peak amplitude earlier than the fastest responses of a broad thorny cell. These findings reveal a complex and nuanced retinal code during naturalistic stimulation.

**Figure 4.**
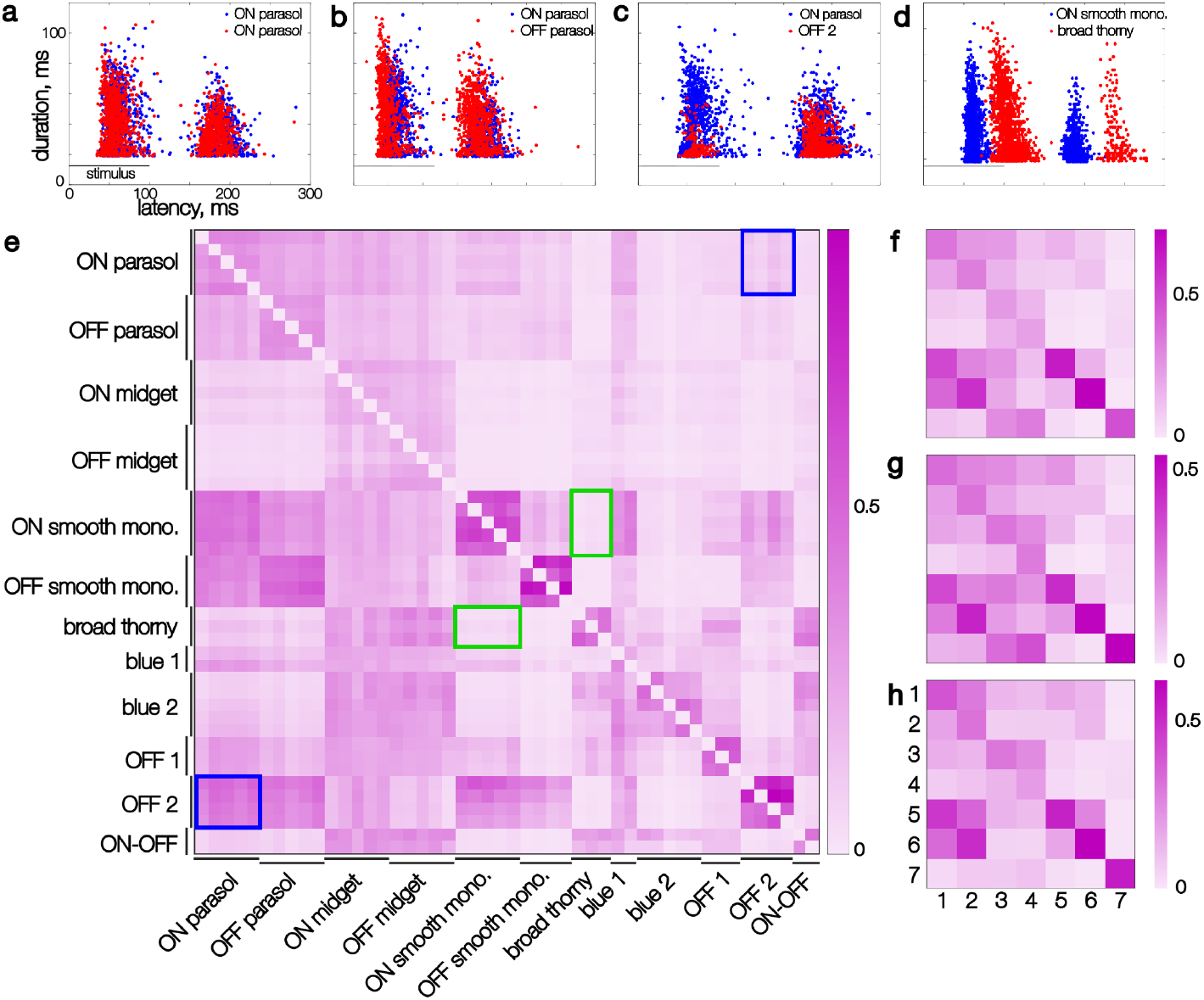
Kinetic features extracted from recorded responses of pairs of simultaneously-recorded cells and their similarity. **(A-D)** Each point represents the latency and duration of the response of one cell to a single flashed image, smoothed in time (see Results). Image presentation period is marked with a horizontal line. **(A)** Two ON parasol cells (red and blue dots, respectively). **(B)** An ON parasol and an OFF parasol cell. **(C)** An ON parasol and an OFF 2 cell. **(D)** An ON smooth monostratified and a broad thorny cell. **(E)** Similarity scores (see Results) for individual cells of 12 distinct types in a single recording, sorted by type. At most 5 cells per type are shown. The rows show the similarity score with the cell listed on the ordinate as the reference cell. Blue rectangles highlight the asymmetrical similarity of ON parasol to OFF 2 cells compared to the reverse. Green rectangles highlight symmetric low scores between broad thorny and ON smooth monostratified cells. **(F)** Similarity scores for the same recording as **(E)**, averaged across each cell type, for 7 identified types (1 through 7: ON parasol, OFF parasol, ON midget, OFF midget, ON smooth monostratified, OFF smooth monostratified, broad thorny). **(G-H)** Same as **(F)**, for two additional datasets.

To quantify similarity in response latency and duration across cell pairs, the inverse of the mean distance to the k-nearest responses was computed (see Methods). This measure is asymmetric: if the responses of cell A form a subset of cell B’s responses (e.g., Fig. 4c), the similarity of A to B will be high, while the similarity of B to A will be lower (compare lower and upper triangles of Fig. 4 matrices).

The response repertoire similarity was conserved for cell pairs composed of specific types within a recording, as revealed in the block-wise structure of the similarity matrix (Fig. 4e). For instance, each broad thorny cell exhibited very low similarity score with each ON smooth monostratified cell, symmetrically (Fig. 4e, green frames). On the other hand, each pair of OFF 2 and ON parasol cells exhibited consistently asymmetrical scores (Fig. 4e, blue frames). The similarity score also reveals internal variability of the response repertoire: cells with tighter distribution of response parameters show higher scores within-type (e.g. within-type pairs of ON and OFF smooth monostratified and broad thorny cells score higher than within-type pairs of parasol, midget, and blue 1 and 2 cells). The type-specific response repertoire was also robust to variability between animals and experiments, as revealed in the consistent pattern of the per-type similarity matrix across experiments for cells of known types (ON and OFF parasol, midget, smooth monostratified, and broad thorny) (Fig. 4f-h). Thus, the well-known distinctive properties of *average* responses of cells of different types also applies to the entire *repertoire* of their responses, within and across experimental preparations.

## Discussion

Large-scale recordings from diverse types of macaque RGCs showed that both the mean response and the full repertoire of responses to natural images are highly stereotyped within each cell type and distinctive in different cell types, including the less-understood low-density cell types. A few cell types exhibited very similar response repertoires and averages to one another, but most types differed substantially from others, in a consistent manner across retinas. Thus, each RGC type exhibits an intrinsic functional signature that is maintained under the complex conditions of naturalistic stimulation.

The distinctions observed have practical implications for understanding cell type diversity in the retina, because responses to natural images highlighted distinctions between types (Fig. 3a) that were not as readily visible with white noise stimulation (Fig. 3b), the standard method for cell type identification in large-scale recordings ^13–15^. A possible reason for this is that naturalistic stimulation engages nonlinear processing mechanisms ^5–7^ that generate richer temporal patterns of response. On a practical level, however, responses to natural images are generally more susceptible to spike-sorting artifacts due to highly correlated stimuli and responses, which could limit the practical utility of this approach. Synthetic stimuli (e.g. ^7^) could potentially leverage some of the advantages of natural images without compromising spike sorting as much, an area for further exploration.

The striking distinctions in response repertoires observed during natural stimulation suggest the possibility that downstream mechanisms in the brain could exploit these patterns during development for cell type specific refinement of synaptic contacts. Many studies of visual system development focus on the molecular mechanisms that contribute to specific connectivity in retinal targets ^2,18,19^. However, a large body of work also points to the importance of visually-driven activity in segregating distinct retinal inputs to central structures ^20,21^. Although a dominant theory is that correlated firing over space is a driving factor ^22–24^, the very distinctive patterns of activity in different RGC types could also be a cue for refinement of retinal inputs to downstream structures, and the present results show that there is enough distinction in response repertoires to potentially support such a mechanism.

Some of the distinctions between cell types were consistent with what would be predicted based on previous studies using simpler non-naturalistic stimuli such as white noise, gratings, and contrast steps ^3,5,6,25^. Specifically, midget RGCs displayed more sustained responses than parasol RGCs ^4,25^, ON and OFF RGCs of morphologically matched types (midget, parasol, smooth bistratified) had very similar response repertoires and mean responses, and OFF RGCs tended to have slightly shorter response latencies than ON RGCs of morphologically matched types ^16^. Thus, some of the major cell type distinctions seen in earlier studies are applicable to natural vision.

Some distinctions between cell types, particularly the less-studied types, have not been reported previously, and could potentially be important for understanding their role in natural visual signaling. The response latencies of smooth cells varied little across images compared to other cell types (Fig. 4), including parasol cells, an invariance that could support a role in signaling the timing of stimuli to the brain. Some less-studied RGC types exhibited consistent response durations across different images, unlike midget and parasol cells which had response durations that varied strongly with image content (Fig. 4), and several cell types exhibited more transient responses than parasol cells (Fig. 2). The mechanisms for these distinctions in response kinetics across cell types are not known. In principle, such differences could arise from different spike generation mechanisms ^26^, synaptic input properties ^27^ or strong inhibitory input from amacrine cells (e.g. ^28,29^).

Two primary limitations of the present work could be important for its interpretation. First, the stimulus set consisted of grayscale flashed natural images, which may not fully engage the dynamic response properties of certain RGC types. Second, the use of only two kinetic features (latency and duration) to analyze cell type distinctions may not reveal the full diversity of responses. Future studies incorporating stimuli with color, object motion, optic flow, as well as analysis of a broader range of kinetic features will be important to understand the full range of RGC response properties across cell types in natural viewing conditions.

## Methods

### Tissue Preparation

Retinas were obtained from macaque monkeys following terminal procedures conducted in compliance with Stanford Institutional Animal Care and Use Committee guidelines. Following enucleation, the eyes were hemisected and the vitreous was removed. Small retinal segments (approximately 2 × 3 mm) with the retinal pigment epithelium attached were dissected from regions 8-10 mm from the fovea, after which the choroid was trimmed to optimize tissue oxygenation.

### Multi-electrode Array Recordings and Spike Sorting

Retinal ganglion cell (RGC) activity was recorded using custom multi-electrode arrays (MEAs) with 512 electrodes arranged in a 16 × 32 isosceles triangular grid with a 60 µm separation between rows and between electrodes in a row, covering roughly 1 × 2 mm. The retina was mounted ganglion cell side down onto the MEA and secured with a permeable membrane. During recordings, the tissue was continuously perfused with oxygenated Ames’ solution (Sigma, St. Louis, MO) maintained at 31-33°C. Voltage signals were band-pass filtered, amplified, and digitized at 20 kHz using custom electronics (Litke2004). Spike sorting was performed with Kilosort2 ^30^, and only cells meeting rigorous quality criteria (no refractory period violations, distinct electrical images, and consistent receptive field properties across cells of each identified type) were included in the analysis.

### Visual Stimulation

Two types of visual stimuli were presented on a computer display and focused onto the photoreceptor layer. The display intensity produced on average 800-2200, 800-2200, and 400-900 photoisomerizations per second for the L, M, and S cones respectively. For cell classification and receptive field mapping, a series of 4-8 white noise stimuli (flickering checkerboards) was presented, each lasting 30-60 minutes and with differing pixel size and refresh time. In these stimuli, the contrast of each pixel was selected randomly and independently over space and from a binary distribution at each refresh. In some cases the three display primaries modulated independently, in other cases the three primaries were yoked. For naturalistic stimulation, grayscale images from ImageNet dataset ^31^ were used. Each image was presented for 100 ms, followed by 400 ms of a uniform gray background at mean luminance, a design intended to isolate individual responses and minimize adaptation. A total of 10,000 unique images were shown, interleaved with a block of 150 repeated images after every 1,000 unique images to monitor recording stability.

### White noise analysis

The responses to white noise were analyzed as described elsewhere ^13–15^. Briefly, the spike-triggered average (STA) from the white noise stimulus was used to extract spatial and temporal response properties, facilitating cell type classification based on time course, inter-spike interval distributions, and mosaic organization, as described in ^14^. Cells were then categorized into known types (e.g., ON/OFF parasol, midget, smooth monostratified types, and broad thorny type ^14^ and other putative types based on similarity of these parameters and the mosaic organization of receptive fields.

To compute the time courses, significant pixels in the STA were identified as follows. First, the dominant STA frame was determined by selecting the frame with the maximal absolute pixel intensity. Within that frame, pixels were considered significant if their intensity exceeded four times the robust standard deviation calculated across all pixels. These significant pixels were then grouped by polarity (sign), and their spatial average was computed for each STA frame separately for each group, resulting in two time courses for each display primary. For the blue and green channels, the time course corresponding to the dominant polarity (i.e., with the highest absolute amplitude) was selected.

An elliptical fit of the spatial receptive field was obtained by fitting a 2D Gaussian to the STA frame with the maximum signal amplitude. The 2-standard deviation contour of this fit is shown in Figure 1 to illustrate the mosaic organization.

### Natural image analysis

For naturalistic stimuli, spike responses were aligned to stimulus onset, smoothed with a Gaussian kernel (σ = 10ms) and averaged over a 500 ms window to obtain the evoked mean firing rate for each cell. To capture the full repertoire of responses, two temporal features for each response were computed: latency (defined as the time to peak amplitude) and duration (defined as the time from the peak until the response decayed to 25% of its maximum). These features were calculated for a random subset of 1500 responses per cell, after excluding trials with zero or single spikes, to ensure only stimulus-driven responses were analyzed. The results were robust to resampling and to varying the subset of selected trials between 500 and 3000 images (across cells, the total number of trials after filtering for the number of spikes varied between 2000 and 9500). Alternate definitions of latency (the time at which half the spikes in the trial were recorded) and duration (full width at half-height for the largest peak) yielded results similar to those obtained with the original definitions. The difference between metrics after normalization was similar in magnitude to the difference across resamplings of 1500 trials. The distance between the response repertoires of two cells was quantified by *(latency*^*2*^*+duration*^*2*^)^*1/2*^. For each response from cell 1, the k smallest distances (typically k = 10) to responses from cell 2 were identified, and the inverse of their average was taken as a similarity score.

### Two-dimensional representation of cell properties

To visualize cell type clusters (Fig. 3), principal components analysis was applied to several groups of parameters: 1) normalized time course for the blue and green display primary (Fig. 1, column 3) concatenated with the normalized interspike interval distribution obtained during white noise stimulation (Fig. 1, column 4); 2) normalized average response time courses (Fig. 2b) for natural image stimuli, 3) a concatenation of 1 and 2. All normalizations were L-2 norm. The t-distributed Stochastic Neighbor Embedding ^17^ was used on the first 8 principal components to project the data into two dimensions.

## Acknowledgements

this work was supported by NIH grants NEI R01-EY029247, R01-EY033870, R01-EY021271, R01-EY032900, and Research to Prevent Blindness Stein Innovation Award (EJC). We thank Eric Wu, Jillian Desnoyer, and Sam Cooler for technical assistance; T. Moore, S. Moriarty, the UC Davis Primate Center for providing access to macaque retinas; Fred Rieke, Michael Manookin, and Greg Field for the insightful discussions, suggestions, and feedback.

## Author contributions

A.K. and E.J.C. conceived the work and designed the multi-electrode array experiments; A.K., C.R., N.B., and A.G. performed multi-electrode array experiments; A.K., A.G., and E.J.C. analyzed the data; A.S. and A.L. provided multi-electrode array technology; A.K. and E.J.C. wrote the manuscript.

